# Same-sex sexual behaviours in a wasp despite sex recognition cues: the broad mating filter hypothesis

**DOI:** 10.1101/2025.11.20.689450

**Authors:** Blandine Charrat, Guillaume Meiffren, Dominique Allainé, Isabelle Amat, Emmanuel Desouhant

## Abstract

Same-sex sexual behaviour (SSB) in animals, once considered an evolutionary dead end, is surprisingly widespread, particularly among male insects. These behaviours may arise from either adaptive or non-adaptive processes. Over shorter timescales, SSB can occur when courtship costs are low or when missing a mating opportunity is costly, leading to a reduced behavioural threshold for courtship—a “broad mating filter,” as predicted by threshold acceptance theory. We investigated SSB in males of the parasitoid wasp *Venturia canescens* (Hymenoptera: Ichneumonidae), a species where encounters are infrequent and male courtship is energetically not costly. We hypothesised that males exhibit a broad mating filter, accepting conspecifics for courtship regardless of sex. First, behavioural assays assessed whether SSB correlated with higher reproductive success. Second, we compared cuticular hydrocarbon (CHC) profiles between sexes, as these compounds may serve as cues for sex recognition. Our chemical analyses revealed clear quantitative CHC differences between males and females, indicating potential for sex recognition. Nonetheless, 18% of males displayed SSB, and these behaviours had no detectable effect on either courting or mating success. Moreover, males receiving SSB did not experience any reproductive advantage. Our findings support the hypothesis of a broad mating filter in *V. canescens* males. In this species, where conspecific encounters are rare, SSB appears to arise from reduced discrimination rather than adaptive benefits, consistent with a low-cost courtship strategy and high opportunity costs of missed mating events.

## Introduction

Same-sex sexual behaviours (SSB) are defined as reproductive behaviour between individuals of the same sex that are generally observed between individuals of opposite sex, such as courtship or mating (Bailey & Zuk, 2009). They are usually considered as misdirected behaviours because they are directed towards the same sex and usually observed in the context of reproduction such as courtship or mounting behaviours. They thus have long been considered as evolutionary dead ends (Sommer & Vasey, 2006). However, their ubiquity in all animal taxa, especially in male insects, is a paradox from an evolutionary point of view, and has stimulated much conceptual and experimental work (Monk et al., 2019; Scharf & Martin, 2013). Two categories of hypotheses have emerged to explain the taxonomically widespread maintenance of these SSB (Bailey & Zuk, 2009; Boutin et al., 2016). On the first hand, adaptive hypotheses support that SSB have indirect reproductive benefits, SSB being a way to learn sexual behaviours (learning hypothesis (Lerch & Servedio, 2025; McRobert & Tompkins, 1988; Sommer & Vasey, 2006)) or to show individual’s competitivity (competitivity hypothesis (Bailey & Zuk, 2009)). On the other hand, non-adaptive hypotheses suppose that SSB have no fitness benefit. The main non-adaptive hypothesis is the sex misidentification hypothesis (Scharf & Martin, 2013); it assumes that individuals fail to recognize or misrecognize the sex of the conspecifics they encounter.

The maintenance of SSB from an evolutionary point of view remains a paradox. However, at a shorter timeframe than the evolutionary one, SSB occurrence in animals can be explained by the threshold acceptance theory (Clutton-Brock, 1984). In ecological behaviour context, this framework suggests that behaviours need a certain level (i.e. threshold) of stimuli (which may be multiple) to be triggered. According to this theory, when a male encounter a conspecific, its choosiness should result from the relative costs of acceptance and rejection errors. In a species where males do not suffer SSB, for example in terms of injury or energy loss, or where high costs are associated with losing a mating opportunity (i.e. encounters with conspecifics are rare), it is expected that the choosiness is low. This low level of choosiness was qualified as a ‘broad mating filter’ by Richardson and Zuk (2023). In contrast species where individuals have a very high acceptance threshold, which may be linked to high costs associated with erroneous courtships or a high probability of encountering suitable sexual partners, have a ‘narrow mating filter’. Different degrees of discrimination and thus different sizes of the mating filter should result from spatial and temporal variations in environmental conditions affecting the relative costs of SSB versus the wrongful rejection of a female. For example, in the cricket *Teleogryllus oceanicus*, males courted other males but also not sexually mature males and females even though they seem to be able to distinguish the sex of conspecifics encountered (Richardson & Zuk, 2023). The authors concluded that males have a broad mating filter, i.e. a low acceptance threshold when courting. More generally, individuals with broad mating filter expressing SSB could benefit from a higher reproductive success linked to their higher probability to court or mate when encountering a conspecific (regardless of its sex), even if sex cues exist in this species.

Different cues can be used for sex recognition in insects. For example, morphological cues can be employed (Bland, 1991), as in the beetle *Diaprepes abbreviatus*, where males engage in SSB towards males of female-like size, using body size as proxy sex recognition (Harari et al., 2000). Literature has shown that chemical cues with sex-specific pheromones (i.e. long-range compounds, see Wyatt, 2014; Yew & Chung, 2015) or cuticular hydrocarbons profiles (CHC profiles, i.e. short-range compounds) are often involved in sex recognition. CHC are long-chain hydrocarbons present on insect cuticle (Blomquist & Bagnères, 2010). CHC profiles carried by individuals is a powerful means of communication due to their wide range of variation (in number of different compounds and/or quantity of each compound). Thus, CHC are relevant source of information about sex (Blomquist & Bagnères, 2010; Hare et al., 2022; Thomas & Simmons, 2008), reproductive status (Hora et al., 2007) or age (Kuo et al., 2012; Nieberding et al., 2008). However, because CHC profiles may be plastic, chemical information they provide may lead to sex misidentification when, for example, young males have a female-like profile as in the parasitic wasp *Lariophagus distinguendus*. CHC profiles between young males and females being similar, these young individuals are not recognized as males and receive SSB from older males (Ruther & Steiner, 2008). Such plasticity also occurs in wild bee, *Osmia cornuta* and *O. bicornis*, where males begin to produce specific CHC as SSB-inhibiting pheromones within 3 days of emergence (Seidelmann, 2023).

This study aims at investigating how and why SSB occur between males of the parasitoid wasp *Venturia canescens*. It is a suitable insect model to approach the mating filter concept. Courtship displays by polygynous males to females are not costly, at least in terms of male longevity (Charrat, 2023) or injury. Moreover, encounters in the field, i.e. potential mating opportunities with virgin females (monandrous females), are not frequent (Collet et al., 2020). Consequently, the cost of losing a mating opportunity by not expressing a courtship should be higher than the cost of expressing a misdirected courtship to males (i.e. SSB). Finally, sex recognition may be mediated by CHC as the female wasps use this information to recognize siblings during mate choice (Chuine, 2014; Collet et al., 2020), probably by comparing their own CHC profile with that of the encountered conspecific (“self-phenotype matching” hypothesis (Lacy & Sherman, 1983; Metzger et al., 2010)).

Using an integrative approach combining behavioural experiments and chemical analyses of CHC profiles, we aimed at explaining the occurrence of SSB in males *Venturia canescens*. First, we tested if there was a link between the expression or the reception of SSB, and the probability of courting and mating when encountering a receptive female. Second, we assessed whether CHC could be a cue for sex recognition. As in this species courtships are not costly and conspecific encounters are rare events, we predicted that *V. canescens* males have broad mating filters. Then, males should not consider the sex recognition cues, leading to SSB expression without benefit in terms of reproductive success.

## Material & Methods

### Biological model and experimental conditions

*Venturia canescens* (Hymenoptera: Ichneumonidae) is a parasitoid wasp in which females are monandrous (Collet et al., 2020; Metzger et al., 2010) and males are polygynous (Charrat et al., 2023). Females lay eggs in host lepidopteran larvae (Driessen & Bernstein, 1999; Salt, 1976). Under laboratory conditions, we used *Ephestia kuehniella* Zeller (Lepidoptera; Pyralidae) larvae as hosts, fed with organic wheat semolina. Males court females using a stereotyped behavioural sequence of 5 steps: Localisation, Orientation, Positioning, Mounting and Mating (van Santen & Schneider, 2002). We defined SSB in males by the sequence of at least the first three steps, we never observed mating between two males. We put the threshold at the first three steps because we observed when a male engaged in a courtship, it never expresses less than those three steps (B.C. pers. obs.). All insect cultures were maintained under constant laboratory conditions: 24 ± 1°C, 50 ± 10% relative humidity, DL 12:12. Wasps naturally feed on nectar or honeydew (Casas et al., 2003; Desouhant et al., 2010), so they were fed with 50% water diluted honey in the laboratory. The wasps came from strains established from numerous individuals captured in the field during the summers 2017 and 2018 near Valence in an organic orchard (Drôme, France).

All experiments were conducted between 10am and 4pm, under the same laboratory conditions as rearing. During the observation we used light produced by two daylight bulbs (60W each). Statistical analyses were performed with R version 4.5.1 (R Core Team, 2020).

### Experience 1: SSB frequency and link with mating probability

We first sought to quantify the frequency of SSB in *V. canescens* males, and then to test whether the expression or reception of SSB subsequently influenced male mating probability. To achieve these aims, we observed and quantified SSB in different pairs of males and then allowed males with contrasting expressions of SSB to mate.

One day before the test, virgin males and females from different culture boxes (strains 2017 and 2018) were kept at emergence and individually placed in a plastic tube with honey *ad libitum*. In order to distinguish males within a given pair, each was anesthetized with CO2 (ca 5 minutes) and identified by a dot of blue or orange water-paint on the thorax. This procedure did not affect reproductive behaviours of *V. canescens* (Fauvergue et al., 2015). The two males forming a pair were randomly chosen from the 2 strains (one from 2017 and one from 2018 to avoid having related males in a pair) and with different dot colours.

The morning of the test, at least two hours before the start of the observation, food was removed to standardize the feeding status of males. They were placed in a clean tube. This allowed to prevent the presence of any chemical marks deposited by males when they were stored (Chuine et al., 2015). Up to 10 pairs of males were observed and filmed (Raspberry Camera HQ) simultaneously, for 90 minutes. At the end of the observation period, males were returned to their individual tubes. The number of SSB expressed by each male was quantified by video analysis using the software BORIS (Friard & Gamba, 2016). 133 pairs of males were observed.

In the afternoon, we created two categories of males from those observed in the morning: SSB+, corresponding to males that had expressed at least one SSB, and SSB-, for males that had not expressed any SSB. We then randomly selected males from the two categories (SSB+ and SSB-) and placed them individually in the presence of a day-old virgin female. To do this, the male and female were rapidly anaesthetised with CO2 and placed in a square plastic arena (10^*^10^*^2cm), each confined to one corner of the arena by a barrier system. When they were awake, we opened the barriers and recorded the sexual behaviours of the males for 30 minutes. Presence/absence of courtship and mating and latency to first courtship were recorded. Up to 6 arenas were observed at the same time; each female was used once. 70 males were tested with a female. Among SSB-males, 9 received SSB (i.e. where in pair with SSB+ males), hereafter named Receiver, (“R”) and 26 did not receive SSB (i.e. in a pair where 0 SSB were done), hereafter named Control (“C”). In the following text, the SSB+ males are named “Emitter” (“E”). Four males were removed from the dataset as they were both emitter (E) and receiver (R), leading to a dataset of 66 males (31 males SSB+ and 35 SSB-).

The size of males was estimated using hind left tibia length as proxy (as in Pelosse et al., 2011) by the means of a binocular loupe (3.2 magnification) with a camera (moticam 1000, software Motic Image plus 2.0).

The effect of difference in size between the two males in a pair on the probability of observing SSB in this pair was tested by the mean of a generalized linear model (GLM) with binomial distribution. When females are present, the link between the probability of courting or mating and the status of the male (E, R or C) was tested by the mean of GLM with binomial distribution. The relationship between male status and latency to the first courtship was tested by using a GLM with Gamma distribution. Male size was added as a covariate in each GLM but the additive effect of this variable or its interaction with the other explanatory variables was not significant. Results are presented from the simplest model after a backward procedure (Crawley, 2012).

### Experience 2: Sex recognition mediated by CHC profiles

This experience aimed at characterizing CHC profiles of males and females and testing whether the profile similarity changed with sex. The CHC profile may vary with age (Kuo et al., 2012; Nieberding et al., 2008), but our behavioural experiments on SSB only tested young males that were one day old. To avoid potential age bias in our conclusions about the sex cues carried by the CHC profile, we included older males and females (7 days old) in the analysis as control conditions. Twenty individuals of each age and sex condition were analysed.

Individuals of both sexes were isolated at emergence and individually placed in a plastic tube (diameter = 3 cm, height = 7 cm) with honey *ad libitum*, for 1 day (“Young”) or 7 days (“Old”). After 1 or 7 days, each wasp was individually immersed in a 1.5 mL amber vial containing 400 µL of ethyl acetate for 6 hours to extract chemical compounds. Since we aimed to minimize sample manipulation and avoid evaporation/re-dissolution steps that can introduce variability or compound loss, 400 µL was the most pragmatic choice to ensure direct injection reproducibility. While hexane is widely used in CHC extraction due to its low polarity and limited ability to penetrate internal tissues, a preliminary solvent comparison showed that ethyl acetate consistently allowed the detection of the same number of CHC compounds as hexane, and in several cases, produced stronger signal intensities. Regarding the extraction time, we acknowledge that recent literature tends toward shorter extraction durations (typically <1 h) to limit contamination by internal lipids. However, given the extremely small mass of individual *V. canescens*, we observed that shorter extraction times frequently resulted in weak GC/MS signals and poorly resolved chromatographic profiles. Our 6-hour passive extraction (without vortexing or sonication) was a compromise aiming to maximize surface recovery without mechanical disruption. Importantly, the insects remained morphologically intact post-extraction, suggesting minimal tissue penetration. Each vial was sealed and subjected to analysis by coupled gas chromatography/mass spectrometry (see Supplementary Materials). Chromatograms were processed with MassHunter software (MassHunter Workstation B.07.00). The 20 highest peaks of chromatograms were considered as the major compounds extracted from *V. canescens*. The quantity of each of these 20 compounds from each individual was approximated as the area under the corresponding peak in the chromatogram. Compounds annotation was based on a comparison of experimental versus reference mass spectra of the NIST, CNRS and Wiley databases associated with bibliographic research. In addition, retention index values, calculated from a mixture of reference standards under the same chromatographic conditions, were used to further refine and validate compound annotation. CHCs are long-chain carbon compounds, such as alkanes and alkenes with 21 to 50 carbon atoms (Blomquist & Bagnères, 2010; Blomquist & Ginzel, 2021). We therefore reduced the list of extracted compounds, keeping only the 6 alkanes and 6 alkenes with 21 to 50 carbon atoms.

We then checked whether CHC profiles varied qualitatively and/or quantitatively according to age and/or sex. As the amount of CHC should be proportional to the size of the individual, we used the normalized area of each peak with a clr transformation (*clr*() function from Hotelling package (Curran & Hersh, 2021)) to ensure that each amount was independent of the others for an individual (Aitchison, 1986) (see Supplementary Materials). We then constructed a between-group PCA (BCA) (Dray & Dufour, 2007) with *bca*() function from ade4 package (Dray & Dufour, 2007). BCA is based on the average amount of each compound in each experimental condition (age and sex), thus avoiding potential problems associated with small sample sizes (unlike plsDA) (Doledec & Chessel, 1987, 1989). First, we tested the existence of different groups according to age and sex through a permutation test. Then in order to ensure that the given representation of our data reflects the reality, we used the cross-validation procedure proposed by Thioulouse et al. (Thioulouse et al., 2021) leading to the calculation of the Δ*Ō*ij index. This index characterizes the percentage of difference between representations of experimental and cross-validated data (see Supplementary Materials). The more Δ*Ō*ij is low, the less representations of experimental and cross-validated data differ and the more the representation of the experimental data is credible (41).

We therefore had 4 categories of individuals: young males, young females, old males, old females. We used linear models to test the effect of the category on the quantity of each compound extracted from individuals. As the quantity of each compound was proportional to the size of the individual by using normalized area of each peak with a *clr* transformation (see before), we did not include size in the models. A post hoc Tukey analysis was applied to test for the compound’s quantities between all sex and age modalities, using the *glht*() function from multcomp package (Hothorn et al., 2021).

## Results

### Experience 1: SSB frequency and link with mating probability

Among the 133 pairs of males, we observed at least 1 SSB in 26% of the pairs, representing 18% of the males that expressed SSB. The difference in size between the two males did not influence the probability of observing SSB in pairs of males (χ^2^ = 1.42, df = 1, p = 0.23). In the context of an adaptive explanation for SSB, we would expect a higher probability of mating for males that had previously experienced (as emitter (E) or receiver (R)) these misdirected behaviours compared to control males (C). However, expressing or receiving SSB prior to meeting a female does not affect the likelihood of mating or courtship during the encounter (for courting: respectively 12/26, 13/31 and 3/6 males courted in C, E and R modalities, χ^2^ = 0.46, df = 2, p = 0.79, and for mating probability: respectively 6/26, 9/31 and 0/9 males mated in C, E and R modalities, χ^2^ = 5.3, df = 2, p = 0.07) or the latency to court (respectively 392, 500 and 83 seconds before the first courtship in C, E and R modalities, F = 2.33, df = 1 and 26, p= 0.12).

### Experience 2: Sex recognition mediated by CHC profiles

Males and females had the same qualitative composition of their CHC profiles: indeed, male and female chromatograms were composed of the same 12 peaks corresponding to alkanes or alkenes (Table 1).

**Table 1:**
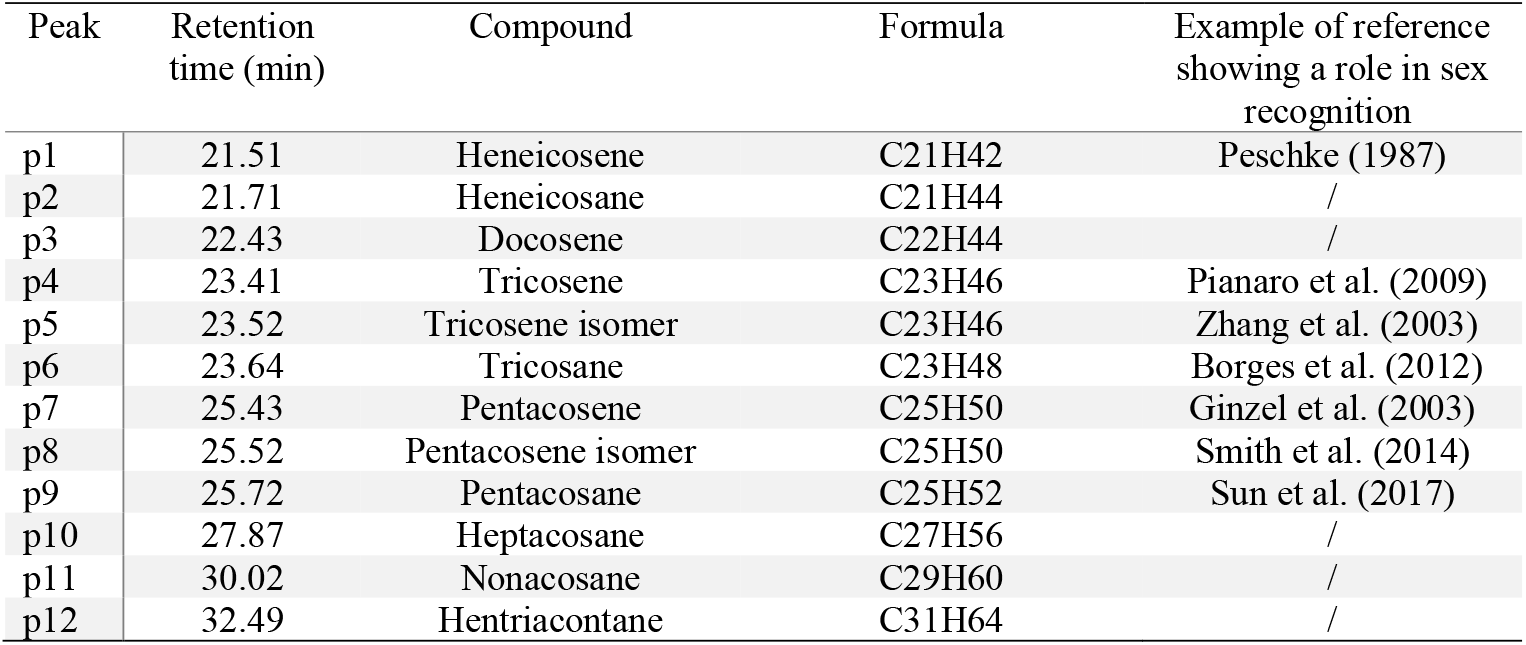
CHC profiles of *Venturia canescens* males and females are composed of 12 main alkanes and alkenes found in the cuticular profiles. Compounds were identified based on a comparison of experimental and reference mass spectra associated with bibliographic research. For each CHCs, we present an example of a reference that shows a role of the compound in sex recognition. / = no identified role in sex recognition in the literature.

| Peak | Retention time (min) | Compound | Formula | Example of reference showing a role in sex recognition |
| --- | --- | --- | --- | --- |
| p1 | 21.51 | Heneicosene | C<sub>21</sub>H<sub>42</sub> | Peschke (1987) |
| p2 | 21.71 | Heneicosane | C<sub>21</sub>H<sub>44</sub> | / |
| p3 | 22.43 | Docosene | C<sub>22</sub>H<sub>44</sub> | / |
| p4 | 23.41 | Tricosene | C<sub>23</sub>H<sub>46</sub> | Pianaro et al. (2009) |
| p5 | 23.52 | Tricosene isomer | C<sub>23</sub>H<sub>46</sub> | Zhang et al. (2003) |
| p6 | 23.64 | Tricosane | C<sub>23</sub>H<sub>48</sub> | Borges et al. (2012) |
| p7 | 25.43 | Pentacosene | C<sub>25</sub>H<sub>50</sub> | Ginzel et al. (2003) |
| p8 | 25.52 | Pentacosene isomer | C<sub>25</sub>H<sub>50</sub> | Smith et al. (2014) |
| p9 | 25.72 | Pentacosane | C<sub>25</sub>H<sub>52</sub> | Sun et al. (2017) |
| p10 | 27.87 | Heptacosane | C<sub>27</sub>H<sub>56</sub> | / |
| p11 | 30.02 | Nonacosane | C<sub>29</sub>H<sub>60</sub> | / |
| p12 | 32.49 | Hentriacontane | C<sub>31</sub>H<sub>64</sub> | / |

However, the BCA revealed that CHC amounts varied according to sex and age, resulting in distinct groups (permutation test p=0.001; Figure 1). The two modalities of the two conditions explained 42% of the total variation. The first axis split individuals according to their sex, with males on the left and females on the right of the representation. The second axis separate individuals of the same sex according to their age, but the signal is less clear for females where old females are confounded with young ones. Overall, young males were closer to old males, and young females were closer to old females. Thus, CHC profiles of males and females were different. Our biological conclusions are supported by the reliability of BCA highlighted by the small value of ΔŌ ij index (0.63%).

**Figure 1:**
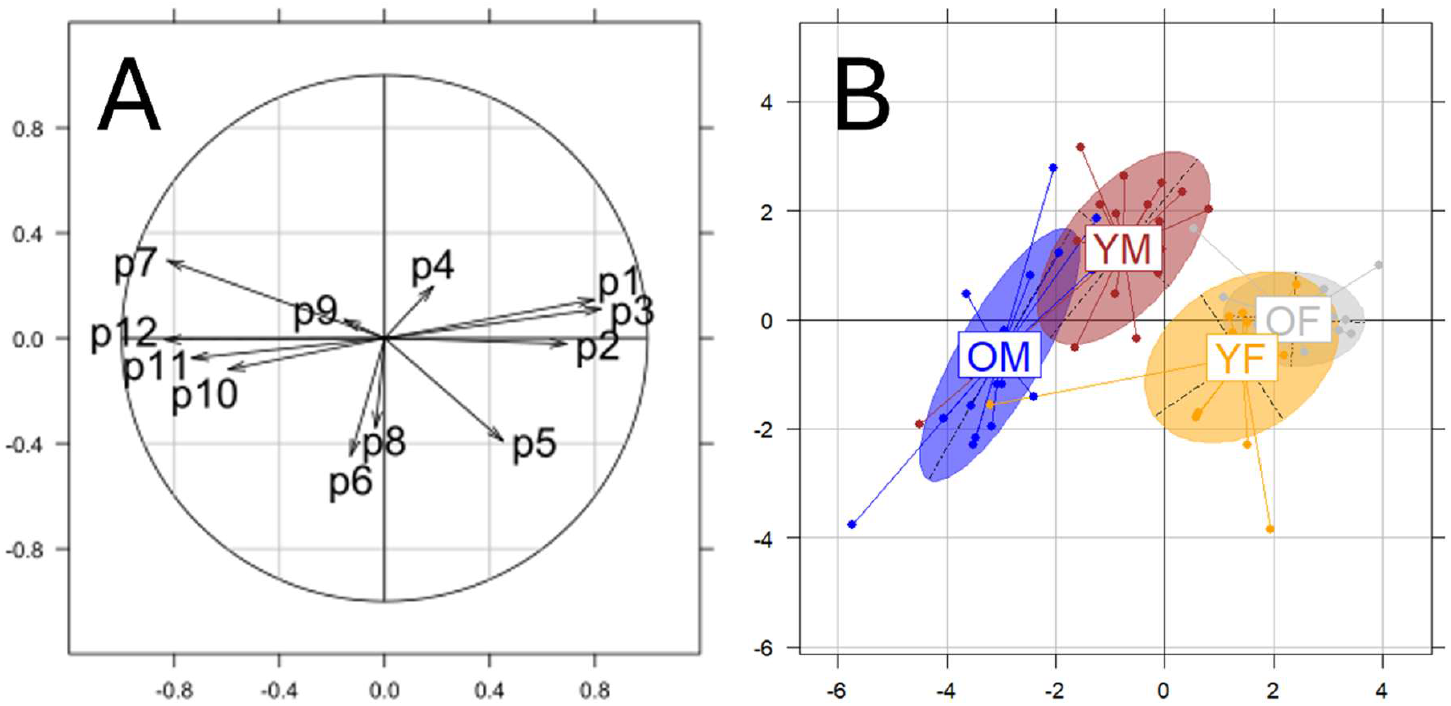
Between-group principal component analysis (BCA) representation of the CHC amounts for the 4 conditions of individuals. A: correlation circle with each peak (named p1 for the first peak, p2 for the second peak, etc, see Table1) corresponding to a compound and B: individual representation in the BCA space. YM = young males, YF = young females, OM = old males, and OF = old females. Each point represents an individual and the label is placed at the barycentre of the point cloud.

Secondly, we compared the quantity of each CHC between the 4 categories of individuals. This revealed that CHC profiles of males and females from both age modalities were also different, with some few exceptions (Table 2). Pentacosane, i.e. compound corresponding to peak 9, was an exception to this result, as it appeared to be in the same quantity in all considered modalities. The quantity in tricosene, i.e. compound corresponding to peak 4, was the same in males and females and only differed between young and old males. This is in accordance with BCA results, where those two compounds correspond to short arrows in the correlation circle (Figure 1), which indicate that they did not strongly contribute to the axis (peaks 4 and 9 contributed 0.8% and 0.5% to axis 1 and 6% and 0.8% to axis 2, respectively). Concerning the effect of age, 9 compounds out of 12 were different between old and young male CHC profiles. However, in females, only one compound (docosene, peak 3) varied between old and young individuals. This confirms what we observed in the BCA (Figure 1) where old females were similar to young ones.

**Table 2:**
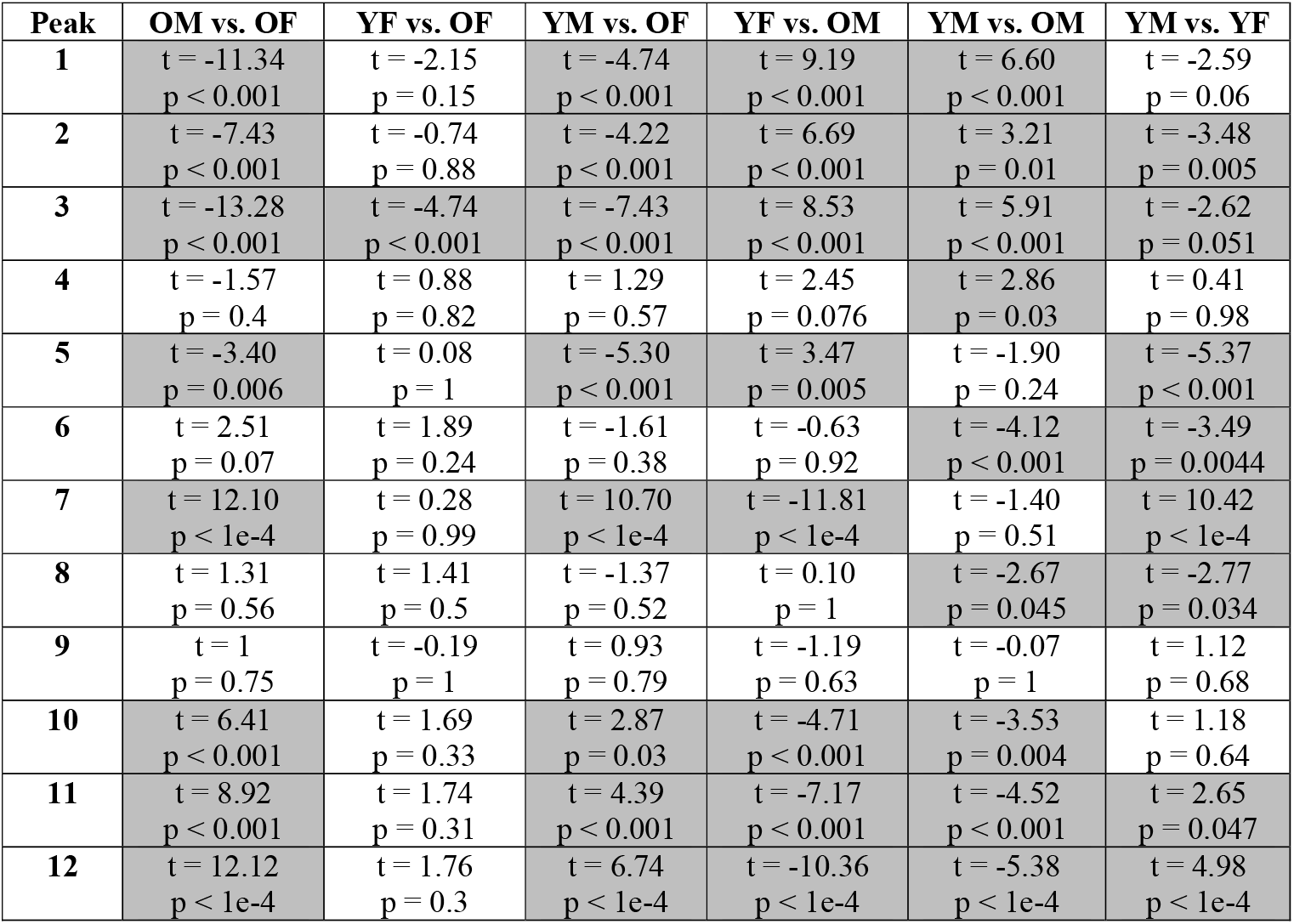
Comparisons of the quantity of each compound between modalities of sex and age. OM = Old Males, OF = Old Females, YM = Young Males, YF = Young Females. Values correspond to t statistic and p-value from the Tukey post hoc analysis. Grey cells indicate significant difference in the quantity of the compound between the two modalities (df = 76).

## Discussion

Same-sex behaviours (SSB) are ubiquitous in animals in particular in insects (Scharf et al., 2013) such as field crickets (*Teleogryllus oceanicus*), damselflies (*Ishnura elegans*) and flies (*Drosophila melanogaster*) where respectively 16%, 17% and 17% of males express SSB (Bailey et al., 2013; Bailey & French, 2012; Van Gossum et al., 2005). In our study, SSB are frequent since 18% of *Venturia canescens* males expressed these misdirected behaviours. In the population we tested, we have demonstrated that SSB is not associated with higher reproductive success in males, at least in terms of the probability of courtship or mating when they encounter a virgin female, and after they have undergone SSB.

The chemical analysis reveals that CHC may be used as cues for sex recognition in *V. canescens*. Indeed, even if the cuticles of males and females carry the same compounds (CHC), the CHC abundance profiles of males are different to those of females, whatever the age modality (except from few compounds, see Table 2). However, it is known that insects can discriminate between CHC that differ in position, number of double bonds or methyl groups, but not between compounds of different chain length (Kather & Martin, 2015; Van Wilgenburg et al., 2010). The variability of CHC profiles between sexes in *V. canescens* concerns, in part, alkanes (peaks 2, 6, 9 to 12, Table 1). As these alkanes are homologous, i.e.they differ only in chain length, it seems that they cannot be distinguished by insects. Thus, sex recognition based on these compounds may be limited in *V. canescens*. Nevertheless, others CHC differing between sexes in *V. canescens* are distinguishable. Alkanes and their corresponding alkenes such as heneicosane and heneicosene (peaks 1 and 2) or tricosane and tricosene (peaks 4 – 6) vary with sex and age. Moreover, some compounds extracted from *V. canescens* individuals have already been shown to be male and female specific in other insect species. For instance, pentacosene is male-specific in *Odontomachus brunneus* ants, and heneicosene is female-specific in *Aleochara curtula* rove beetle (Peschke, 1987; Smith et al., 2014). Those results combined with those of previous study (Metzger et al 2010) show that *V. canescens* use information from CHC profiles for sex recognition and sib-mating avoidance.

Even if cues for sex recognition are present, our experiment strongly suggests that males ignore this information when encountering a conspecific (male or female) and decide to court. The expression of this SSB has no impact on reproductive success for the males. Indeed, we show that SSB expression or reception in males before encountering a virgin female does not influence their mating probability. The opposite observation was made in the water strider *Tenagogerris Euphrosyne* where males exhibiting higher rates of SSB achieve a higher mating success under conditions of intense scramble competition (Han & Brooks, 2015). Moreover, males *V. canescens* that expressed or received SSB (R and E males) and those that did not expressed or received SSB (C males) had the same probability of displaying courtship when encountering a female. These two results therefore argue in favour of a non-adaptive hypothesis in *V. canescens*, and this conclusion is consistent with that drawn from most studies of SSB in insects (Scharf et al., 2013). Additional experiments would be necessary to investigate thoroughly the evolutionary aspects of SSB in *V. canescens*. Nevertheless our results can be interpreted within the conceptual framework of acceptance threshold theory and mating filters framework (Engel et al., 2015; Richardson & Zuk, 2023). In *Venturia canescens*, several arguments are in favour of a broad mating filter. First, the low cost of male courtship (at least in terms of longevity (Charrat et al., 2023)) is a condition favouring the occurrence of error-generating behaviour of males courting another male rather than a female, i.e. SSB. Since courtship is not costly, not displaying it when encountering a conspecific should be detrimental for male lifetime reproductive success, as it increases the risk of losing a mating opportunity but does not impose any costs on the male (e.g. injuries). This is all the more important in *V. canescens* as the encounter rate between sexual partners is low *in natura* (Collet et al., 2020). Therefore, the costs of discrimination may be high in terms of lost opportunities. Furthermore, previous experiments suggest an advantage in the scramble competition for the male that gets closer to a virgin female first (Charrat et al., 2023). If this is the case, males that make decisions quickly (and risk making a mistake by being inaccurate) should have higher reproductive success than slower (but perhaps more accurate) males.

The SSB expression in males may also be state-dependent. Indeed, the probability of expressing SSB varies greatly in *V. canescens*. Most of males do not express SSB and among the 18% of males expressing SSB, the number of SSB observed for a given male ranged from 1 to 7. Thus, every male may not have the same mating filter, that may be dependent on the condition of the individual. In male crickets *Teleogryllus oceanicus*, males in poor condition courted the other males less, suggesting that SSB expression is plastic and potentially costly in terms of energy expenditure (Richardson et al., 2024). In *V. canescens*, it would be interesting to determine if there is a difference between males in SSB expression linked to their energetic resources. It is known that, in this species, lipidic resources (a high sources of energy) decrease with age (Pelosse et al., 2007) without lipogenesis (Casas et al., 2003). Then, old males in poorer condition may change their mating filter and consequently change their probability to court another male. We can make two predictions. First, older males in poorer condition express terminal investment (Clutton-Brock, 1984). So, they would further widen their mating filter, leading to more SSB. This terminal investment is observed in *V. canescens* females when they increase their exploitation of host patch in order to lay more eggs in harsh environmental conditions (Amat et al., 2006). Second, we expect that males in poorer condition conserve their energy and become choosier, reducing their mating filter, as in *T. oceanicus* (Richardson et al., 2024). This would lead to a decrease in courtships towards other males, perhaps using available chemical information about sex, reducing the likelihood of expressing SSB.

Evidence of frequent SSB and the potential plasticity of this behavioural response, that is, whether the occurrence of SSB decreases with males getting older or in poorer condition, opens up relevant avenues of research in the study of sexual behaviours in this species.

## Author contribution

B. Charrat, D. Allainé, I. Amat and E. Desouhant contributed to the study conception and design. Experiments and analysis were performed by B. Charrat. GCMS was designed and conducted by G. Meiffren. All authors read and approved the final manuscript.

