## Supplementary material for "Same-sex sexual behaviours in a wasp despite sex recognition cues: the broad mating filter hypothesis"

---

### Supplementary Materials 1: Detailed protocol and statistical analyses for Experiment 1: Sex recognition mediated by CHC and relation with individual age

#### Compound extraction and GC-MS analysis

Individuals aged 1 (young) or 7 (old) days were individually immersed in 400 µl of ethyl acetate (HiPerSolv CHROMANORM®, VWR Chemicals) for 6 hours in order to extract the chemical compounds making up the cuticle. The resulting solutions were then analysed by GCMS. A Hewlett-Packard 6890N gas chromatography system coupled to an HP 5973 mass spectrometer (Agilent Technologies) and a DB-5MS column (60 metres long, diameter 0.25 mm, film thickness = phase = 0.25 µm; Agilent Technologies) were used. A volume of 2 µL of solution was injected in splitless mode (= all of the flow is directed into the column, allowing the transfer of all the sample for maximum sensitivity) with helium as the carrier gas (at a constant flow rate of 2.3 mL/min), under a temperature in the injector of 300°C. The temperature gradient used was as follows: 90°C for 3 min, then increase to 140°C at 6°C/min, increase to 240°C at 15°C/min, then increase to 310°C at 6°C/min, and finally 310°C for 6 min. The ionisation source was an electron impact at 70 eV and mass spectra were recorded between 35 and 500 m/z. In each condition, 20 individuals were tested.

#### GC-MS data pre-processing

The quantity of each compound in the sample taken for GCMS is represented by the area under each peak of the chromatogram for each individual. The data were first pre-processed with Mass Hunter Qualitative Analysis (MassHunter Workstation Software B.07.00, Agilent Technologies) consisting of a manual peak integration for each chromatogram. This allowed the creation of a data matrix with all the peak areas aligned by retention times. All these peak areas became our set of variables used for the following statistical analysis. The main peaks were identified and numbered (Figure S1). These 20 main peaks were defined as being present in at least 50% of individuals and different from the background. On chromatograms with different retention times, 30 measurements of the area under the curve corresponding to background noise were taken and then peaks with an area greater than the mean of the background noise were considered to be present in the individual.

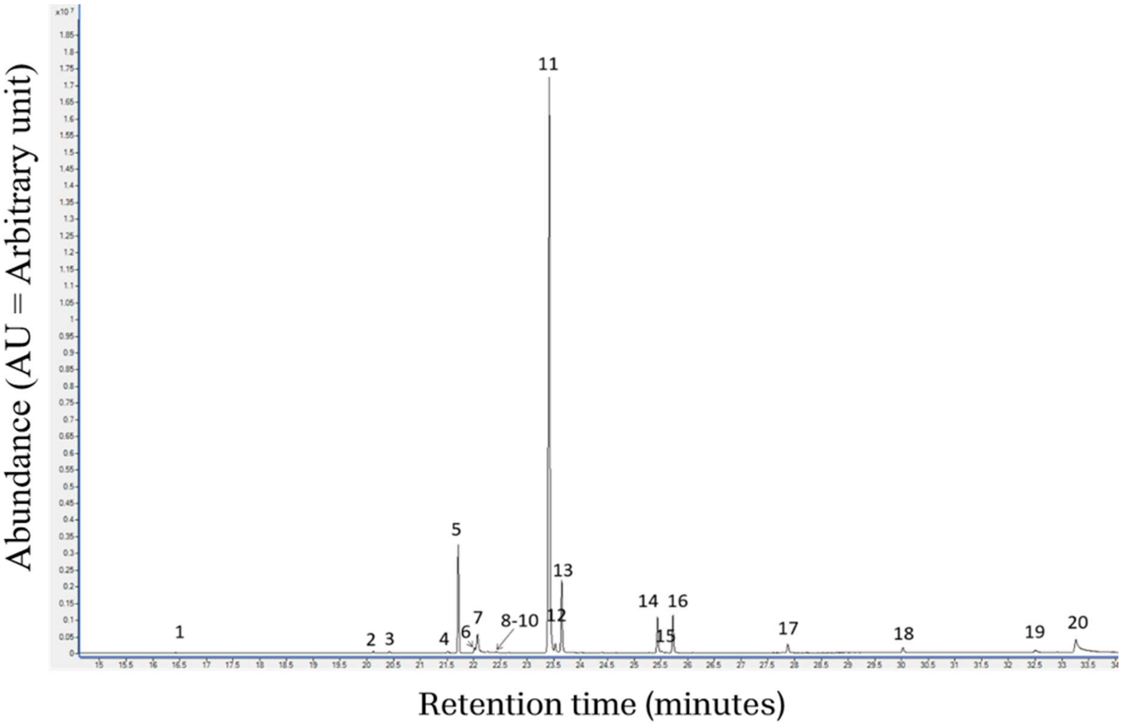


**Figure S1**: Majority peaks selected for analysis of CHC profiles in *V. canescens*.

**GC-MS data annotation**

The nature of the chemical compounds corresponding to the previously identified peaks was determined by comparing the experimental mass spectra (from the chromatograms) by comparison of their retention indices and MS spectra with those reported in the literature (Adams, 2007; Linstrom, 1997) and using Masshunter Qualitative Analysis software (B07.00, Agilent Technologies) matching with standard reference databases (NIST23, Wiley275 and CNRS libraries). During this comparison, an annotation score is assigned (corresponding to the similarity between the experimental spectrum and the most similar reference spectrum), and the higher this score (close to 100), the greater the similarity between the two spectra compared and therefore the more reliable the annotation of the compound. In addition to the score, a retention index (calculated from a mixture of reference standards under the same chromatographic conditions) and a literature search can be used to determine the nature of the compound for each peak. The retention index of each component was calculated relative to a standard mix of n-alkanes (C7-C40, Sigma-Aldrich), analyzed under identical experimental conditions.

**Statistical analysis: BCA**

As the quantity of CHCs is probably proportional to the size of the individual carrying them (as the larger the individual, the larger its cuticle and therefore the greater the CHC layer), so is the quantity of CHCs present in the sample analysed by GCMS. The normalised area of each peak was calculated, corresponding to $A_{ij\_norm} = \frac{A_{ij}}{\sum_{y=1}^{n} A_{iy}}$ with $A_{ij}$ the area of peak j in individual i and $\sum_{y=1}^{n} A_{iy}$ the sum of the areas of the n peaks in individual i. This corresponds to a normalisation of the data. However, the normalised areas obtained in one individual are no longer independent of each other, so a log-transformation must be applied (Aitchison, 1986). The data were therefore transformed using the clr method (clr() function from the Hotelling package (Curran & Hersh, 2021) corresponding to A = $log(\frac{A_{ij\_norm}}{G(A_i)})$ with $A_{ij\_norm}$ the normalised area of peak j in individual i and $G(A_i)$ the geometric mean of the normalised areas in individual i. In order to calculate the geometric mean, 10 were added to each raw value, thus eliminating the 0s (corresponding to an absence of the compound or a quantity too small to be detected) while adding a value small enough in relation to the raw values (ranging from 0 to 85 664 094, median = 473 036.5) not to affect the calculation of the mean (Butterworth et al., 2020; Hervé et al., 2018).

In order to carry out a descriptive analysis of the individual data, an interclass PCA (= BCA) was constructed (ade4 package (Dray & Dufour, 2007)), which maximises the variance between conditions (age and sex) and shows which compounds may characterise each condition studied. The inter-class PCA is based on the average transformed areas for each of the peaks over all the individuals in each condition, which avoids any problems with the number of individuals in the classes. To ensure that the representation obtained corresponds to reality and is not due to a low number of individuals in certain conditions - and therefore to the presence of erroneous groups - an index $\Delta\bar{O}_{ij}$ has been calculated. It is obtained after cross-validation using the permutation procedure proposed by Thioulouse et al. (Thioulouse et al., 2021). The lower this index, the greater the chance that our representation will be close to the cross-validation results and therefore faithful to reality, especially if the percentage of variance explained is itself high.

**Supplementary Materials 2: List of the main compounds extracted from *Venturia canescens***

20 main peaks were extracted and identified in *V. canescens* (Table S1). For CHC profile analysis, we lonely retained the 6 alkanes and the 6 alkenes (compounds corresponding to cuticular hydrocarbons, see main text). Here, we present the BCA analysis based on the 20 main peaks (Figure S2). The eight different chemical compounds of CHC that were extracted are certainly due to the length of time individuals were immersed in the solvent (6h). The peaks 1, 2 and 8 were removed from the dataset for the BCA because they were impurities or we were not sure about their identification.

**Table S1**: 17 main compounds found in the cuticular profiles of *Venturia canescens* males and females. Compounds were identified based on a comparison of experimental and reference mass spectra associated with bibliographic research. The peaks 1, 2 and 8 were removed from the dataset for the BCA because they were impurities or we were not sure about their identification.

| Peak | Retention time (min) | Compound | Formula |
|---|---|---|---|
| p3 | 20.41 | Hexadecanoic acid | C16H32O2 |
| p4 | 21.51 | Heneicosene | C21H42 |
| p5 | 21.71 | Heneicosane | C21H44 |
| p6 | 22/01 | Octadecadienoic acid | C18H32O2 |
| p7 | 22.07 | Octadecenoic acid | C18H34O2 |
| p9 | 22.35 | Octadecenoic acid, ethyl ester | C20H38O2 |
| p10 | 22.43 | Docosene | C22H44 |
| p11 | 23.41 | Tricosene | C23H46 |
| p12 | 23.52 | Tricosene isomer | C23H46 |
| p13 | 23.64 | Tricosane | C23H48 |
| p14 | 25.43 | Pentacosene | C25H50 |
| p15 | 25.52 | Pentacosene isomer | C25H50 |
| p16 | 25.72 | Pentacosane | C25H52 |
| p17 | 27.87 | Heptacosane | C27H56 |
| p18 | 30.02 | Nonacosane | C29H60 |
| p19 | 32.49 | Hentriacontane | C31H64 |
| p20 | 33.24 | Cholesterol | C27H46O |
